# Genomic targets of HOP2 are enriched for features found at recombination hotspots

**DOI:** 10.1101/2023.01.25.525520

**Authors:** Jenya Daradur, Mohamad Kesserwan, Nowlan H. Freese, Ann E. Loraine, C. Daniel Riggs

## Abstract

HOP2 is a conserved protein that plays a positive role in homologous chromosome pairing and a separable role in preventing illegitimate connections between nonhomologous chromosome regions during meiosis. We employed ChIP-seq to discover that Arabidopsis HOP2 binds along the length of all chromosomes, except for centromeric and nucleolar organizer regions, and no binding sites were detected in the organelle genomes. A large number of reads were assigned to the HOP2 locus itself, yet TAIL-PCR and SNP analysis of the aligned sequences indicate that many of these reads originate from the transforming T-DNA, supporting the role of HOP2 in preventing nonhomologous exchanges. The 292 ChIP-seq peaks are largely found in promoter regions and downstream from genes, paralleling the distribution of recombination hotspots, and motif analysis revealed that there are several conserved sequences that are also enriched at crossover sites. We conducted coimmunoprecipitation of HOP2 followed by LC-MS/MS and found enrichment for several proteins, including some histone variants and modifications that are also known to be associated with recombination hotspots. We propose that HOP2 may be directed to chromatin motifs near double strand breaks, where homology checks are proposed to occur.

## Introduction

Meiosis is an evolutionarily conserved, specialized nuclear division process that plays several roles in sexually reproducing organisms (reviewed in Mercier et al. 2015; Wang and Copenhaver 2018). Upon commitment to meiosis, DNA replication takes place and is followed by a relatively protracted prophase, during which unique chromosome behaviors occur. As compaction proceeds, homologous chromosomes move into close alignment and in most organisms, numerous double strand breaks (DSBs) are generated by the SPO11 topoisomerase complex. The free ends are further resected by the MRN complex, generating regions of single stranded DNA (ssDNA), and mutant analyses have identified several proteins that are involved in probing homology and catalyzing the strand invasion (D-loop formation) and repair activities that can lead to reciprocal genetic exchange (reviewed in Crickard and Green 2018; Lam and Keeney 2014). Beginning in leptotene, components of the synaptonemal complex (SC) are deposited along the chromosome axes, and during pachytene, the fully formed SC holds the condensed homologs in precise register and recombination activities are completed. These processes maintain bivalent association and ensure proper homolog segregation at meiosis I, and the reciprocal genetic exchange events contribute to genetic diversity of the progeny. A second meiotic division, in which sister chromatids segregate, results in a reduction in the chromosome number to one half that of the somatic progenitor cells (that is, haploids for diploid organisms). In higher organisms, differentiation takes place to produce sperm and eggs, and upon their union, the diploid state is restored.

The molecular activities of enzymatic complexes that catalyze the events of recombination have been extensively studied in a variety of organisms (Crickard and Green 2018). One of the key, and as yet unanswered questions of meiosis relates to the mechanism by which homology is tested such that precise pairing of homologous chromosomes occurs. Several proteins are known to play roles in the homology search. HOMOLOGOUS PAIRING PROTEIN 2 (HOP2) and MEIOTIC NUCLEAR DIVISION 1 (MND1) form a heterodimer and are instrumental in the modulation of the recombinase/repair proteins DMC1 and RAD51 (Chen et al. 2004; Petukhova et al. 2005; Enomoto et al. 2006; Pezza et al. 2006; Chi et al. 2007; Ploquin et al. 2007; Vignard et al. 2007; Moktan et al. 2014; Zhao et al. 2014; Zhao and Sung 2015; Tsubouchi et al. 2020). Null mutants of *hop2* and *mnd1* exhibit non-homologous chromosome interactions leading to chromatin bridges followed by fragmentation at division, and they display significantly reduced fecundity or meiotic arrest, depending on the organism studied (Leu et al. 1998; Gerton and DeRisi, 2002; Schommer et al. 2003; Petukhova et al. 2003; Zierhut et al. 2004; Kerzendorfer et al. 2006; Panoli et al. 2006; Stronghill et al. 2010; Uanschou et al. 2013; Shi et al. 2019). These studies reveal a positive role for HOP2 in chromosome synapsis and crossing over. Our recent work on wildtype and *hop2-1* haploids in Arabidopsis revealed that HOP2 also has a separable role in preventing non-homologous recombination (Farahani-Tafreshi et al. 2022).

In vitro studies have demonstrated that HOP2 and MND1 are capable of binding to both single and double stranded DNA, can promote strand invasion, and that HOP2 alone can act as a recombinase (Petukhova et al. 2003; Chen et al. 2004; Zierhut et al. 2004; Petukhova et al. 2005; Enomoto et al. 2006; Pezza et al. 2006; Chi et al. 2007; Ploquin et al. 2007; Vignard et al. 2007; Uanschou et al. 2013; Moktan et al. 2014; Pezza et al. 2014; Zhao et al. 2014; Zhao and Sung 2015; Kang et al. 2015, Tsubouchi et al. 2020). Despite this wealth of knowledge about the structure, biochemical properties and protein interactions of HOP2, it is unclear how it exerts its function in probing homology. We sought to better understand whether HOP2 binds to particular DNA sequences and/or to chromatin motifs by conducting both ChIP-seq and mass spectroscopy (MS) of immunoprecipitated HOP2. Here we report that HOP2 binds along the length of all chromosomes except for the centromeric and nucleolar organizer (NOR) regions, and our MS studies reveal possible associations with histone variants and their modifications.

## Results

### HOP2 binds along the length of chromosomes but not to centromere or NOR regions

Several studies have reported the binding of HOP2 and/or the HOP2/MND1 complex to either single or double stranded DNA, or both types of molecules in a variety of organisms (Chen et al. 2004; Pezza et al. 2006; Uanschou et al. 2013; Moktan et al. 2014; Zhao et al. 2014; Kang et al. 2015). Additionally, and depending on the species being investigated, these DNA binding activities can be assigned to regions at the N or C terminus of the protein. In Arabidopsis, HOP2 (At1g13330.1) does not efficiently bind to DNA alone, but does so in the presence of MND1, and the N-terminus of the protein is important for this activity (Uanschou et al. 2013). To our knowledge, all DNA binding studies on HOP2 have employed in vitro assays with either bulk DNAs or nonspecific molecules (e.g. phi X174 DNA). We hypothesized that HOP2 may bind to particular DNA sequences or to specific chromatin motifs, and we therefore designed experiments to test this hypothesis in vivo by chromatin immunoprecipitation and DNA sequencing (ChIP-seq), and by mass spectrometry analyses of immunoprecipitated samples (LC-MS/MS).

We constructed a tagged gene in which a 3X HA epitope is encoded at the carboxyl terminus of HOP2, and transformed *HOP2-1/hop2-1* heterozygous plants. BASTA-resistant T2 transgenic lines were then genotyped to identify tagged lines that are in a *hop2-1* null background (LC831). All developmental stages of the plants were phenotypically wildtype and upon the transition to flowering, LC831 plants were fertile, suggesting that the HOP2pro>HOP2::3XHA construct was able to rescue the *hop2-1* mutant phenotype (Fig. S1).

We conducted ChIP-seq on inflorescences of both LC831 and the parental background line, L*er*. We employed micrococcal nuclease (MNase) digestion to generate chromatin fragments that ranged from mononucleosomes to trinucleosomes (∼150-500bp) and sequenced the DNA from fragments that bound to an anti-HA antibody. Bioinformatic analyses revealed 292 binding sites (q<0.05) distributed along the lengths of all five chromosomes, yet all centromeric (CEN) regions and the nucleolar organizer (NOR) regions on chromosomes 2 and 4 were mostly devoid of binding sites, as were the organelle genomes (Fig. 1).

**Figure 1.**
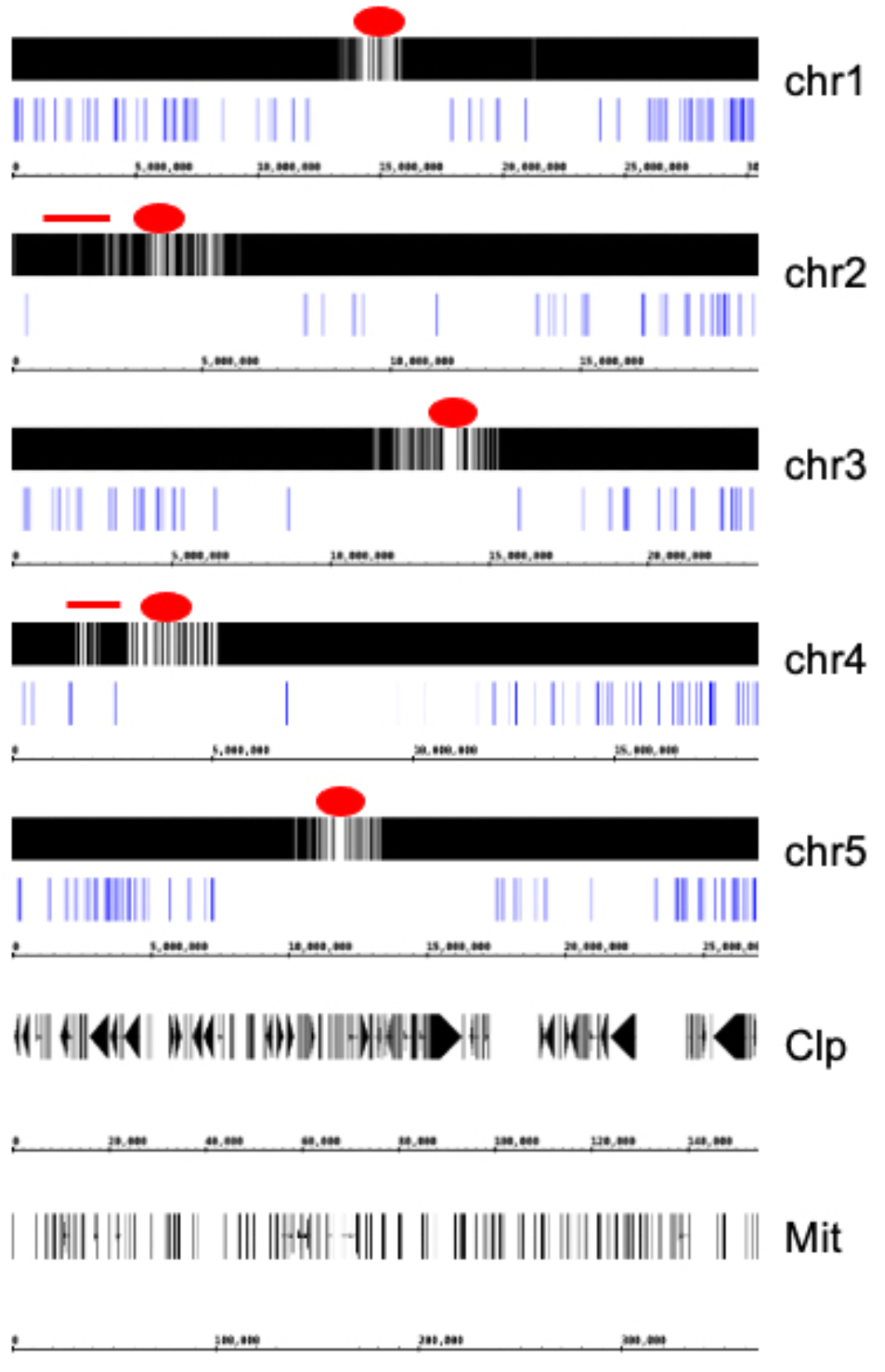
Genome wide binding sites for HOP2. Background signals from L*er* were subtracted from LC831 data and scaled data plotted onto the annotated *A. thaliana* genome using the Integrated Genome Browser. Each of the five chromosomes and the chloroplast (Clp) and mitochondrial (Mit) genomes are shown. The red dots indicate centromeric regions and the red bars on chromosomes 2 and 4 are the NOR loci. The blue lines below each chromosome represent the 292 peaks called by MACS2; no peaks were called for the organelle genomes.

The lack of binding to the NOR regions was surprising, and could be due to loss of these sequences if MNase preferentially cleaves rDNA loci. We used PCR and qPCR to test for the presence of rDNA in MNase digested samples, and observed significant amplification of this DNA even after much of the input DNA had been degraded by MNase (Fig. S2). In contrast, the HOP2 locus was progressively degraded during the time course, reducing the yield of PCR products. Thus, the lack of HOP2 binding to NOR regions is not due to loss of the rDNA sequences during sample preparation, and argues that HOP2 may not exert its functions on this subdomain of chromatin.

Although we did not test for the presence of centromeric sequences in our bound fractions by qPCR, the were a significant number of reads for sequences around the centromeres in both the experimental and control samples, though no enrichment was observed (Fig. S3). We conjecture that centromeric chromatin also is not subject to HOP2 action.

To validate the ChIP-seq results we chose five high scoring regions for HOP2 binding and conducted qPCR on input and bound fractions to ascertain whether these sequences are enriched in the bound fraction. Most of these high scoring regions were slightly enriched, to levels similar to those observed by ChIP-seq (Fig. 2).

**Figure 2.**
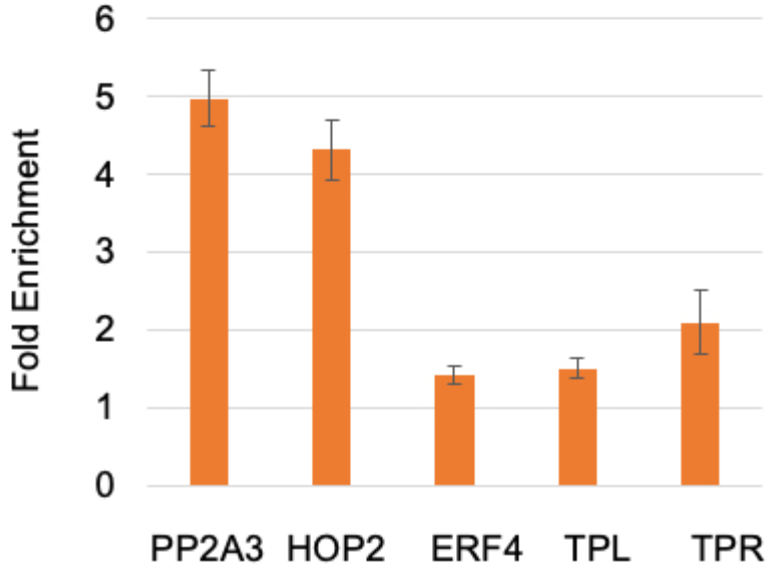
Validation of ChIP-seq by qPCR. DNA from LC831 and Ler chromatin bound to anti-HA beads was subjected to qPCR using primers that bracket enriched loci identified by ChIP-seq. The data was normalized to values for the RTM3 gene, which was chosen based on a similar number of read sequences in both samples and its distance from the nearest peak (>100kb).

Surprisingly, the highest scoring peak identified by ChIP-seq was at the HOP2 locus itself, and in contrast to the average length of 715bp for the 292 peaks, the HOP2 peak encompassed 6.6kb, covering HOP2 and the adjacent PP2A3 coding sequences. For this region, we used two primer sets separated by 3.3kb and both primer sets robustly validated enrichment in the bound fraction. The first primer set amplifies a region near the 3’ end of the PP2A3 gene, while the second primer set is located in the 660bp region that separates the start codons of the two genes.

This curious result led us to examine the sequence reads of this region in more detail. We found that the region of enrichment was situated between the forward and reverse binding sites for PCR primers used to amplify genomic DNA in generating the HA-tagged HOP2 construct (LC831; Fig. 3a). This coincidence suggested that the immunoprecipated sequences aligning to the *HOP2/PP2A3* locus originated from the transgene insertion site (which by TAIL-PCR and sequencing was found to be on chromosome 3, data not shown), and not the native *HOP2/PP2A3* locus. To test this hypothesis, we examined the aligned sequences, using single-nucleotide polymorphisms (SNPs) to distinguish transgene sequences (derived from Col-0) from native HOP2 sequences (the *hop2-1* mutant, which is in a L*er* background). Using Integrated Genome Browser to inspect the region, we identified six SNPs within the region. As expected, all sequences from the L*er* control sample contained the L*er* allele, while nearly all sequences from the LC831 experimental samples contained the Col-0 allele, indicating they arose from the transgene.

**Figure 3.**
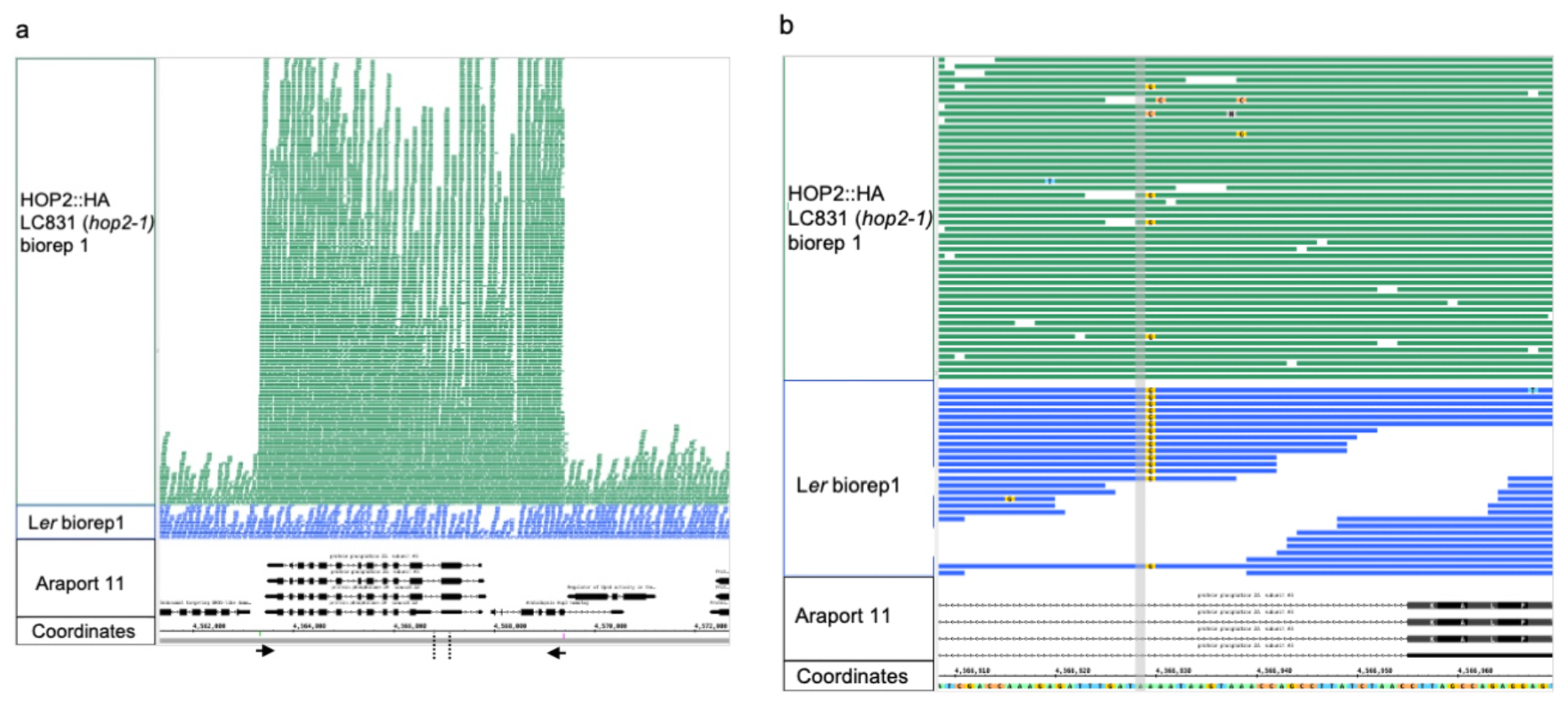
Integrated Genome Browser view of immunoprecipitated sequences aligning to the *HOP2/PP2A3* locus. **A**. The *HOP2/PP2A3* region is enriched for sequences that match the LC831 transgene (green bars) versus the endogenous HOP2/PP2A3 sequences in the L*er* control (blue bars). Arrows at the bottom indicate the location of PCR primers used to create the HA-tagged HOP2 transgene from Col-0 DNA. The vertical dashed lines at the bottom delineate the small region that is expanded in panel b. **B**. A zoomed-in view showing one SNP that distinguishes sequences derived from L*er* (endogenous locus) versus Col-0 (LC831 locus). In this view, letters superimposed on blue and green bars indicate differences between aligned sequences and the reference Col-0 genome sequence. All aligned sequences in the L*er* control sample contain the L*er* allele, whereas most of the sequences in the LC831 experimental samples contain the Col-0 allele. The A/G SNP is highlighted in orange.

However, outside the amplified region, sequences from both samples contained the L*er* allele, indicating they arose from the native loci. While the *HOP2/PP2A3* locus could serve as a pairing center, these results indicate that HOP2 actively discriminates between endogenous and transgene sequences, possibly by identifying regions that are not completely homologous (see the Discussion).

### HOP2 binding sites exhibit commonalities with recombination hotspots

Using ChIPpeakAnno (Zhu et al. 2010), we mapped the location of HOP2 binding sites to discrete chromatin domains and found that the majority of peaks reside in gene regulatory regions such as promoters (75%) and UTRs (44%), with peaks approximately equally divided between genes and intergenic regions (Fig. 4a). At the promoter level, 36% of the peaks center around the transcription start sites, suggesting that HOP2 preferentially binds to open chromatin (Fig. 4b).

**Figure 4.**
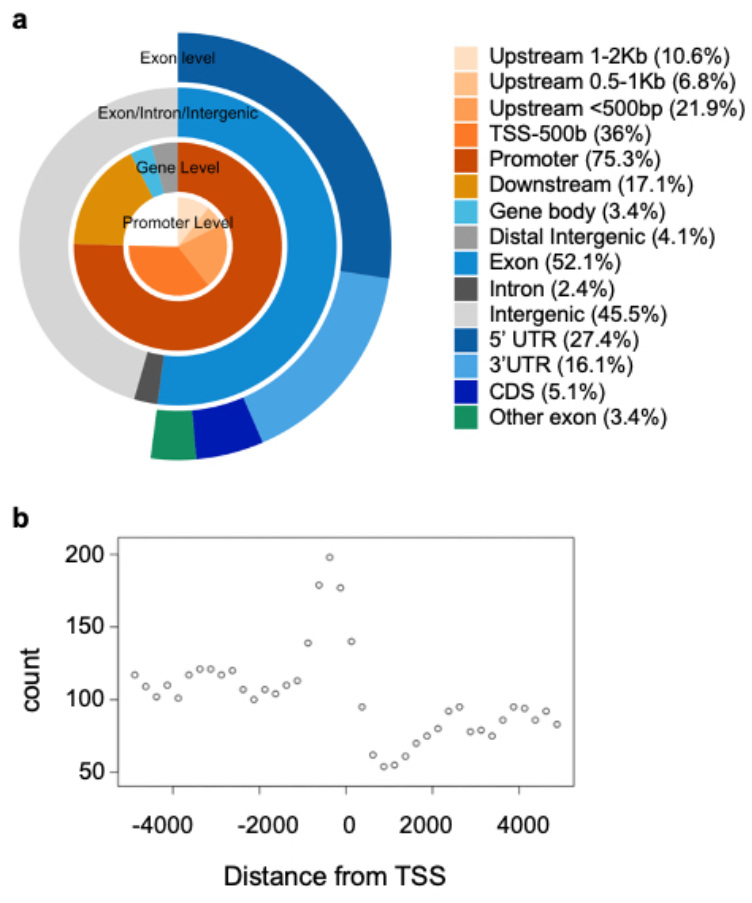
Distribution of HOP2 peaks in annotated genome regions. **a**. The 292 HOP2 peaks were assigned to functional regions of the genome using ChIPpeakAnno genomic Element Distribution v.3.26.0 **b**. ChIPpeakAnno binOverFeature was employed to plot the distance of peaks from transcription start sites (TSS). The number of bins was set to 20 upstream and 20 downstream.

We utilized MEME-ChIP (Machanick and Bailey 2011), to search for conserved sequence motifs that could underpin HOP2 binding to chromatin. MEME identified four motifs with E values less than 2.9e-002 (Fig. 5). An A-rich and a CT-rich motif were found 87 times (E value= 3.2e-016) and 57 times (E value = 1.3e-045), respectively, and there were fewer occurrences of two G rich motifs with higher E values. Three of the four motifs are similar to the binding sites of known transcription factors from the BBR/BPC, REM, and CAMTA families, suggesting that HOP2 might play a role in modulating transcription. The analysis was repeated with STREME [Simple, Thorough, Rapid, Enriched Motif Elicitation (Bailey 2020)], which identified similar motifs, all having p-values less than 0.039 (Fig. 5).

**Figure 5.**
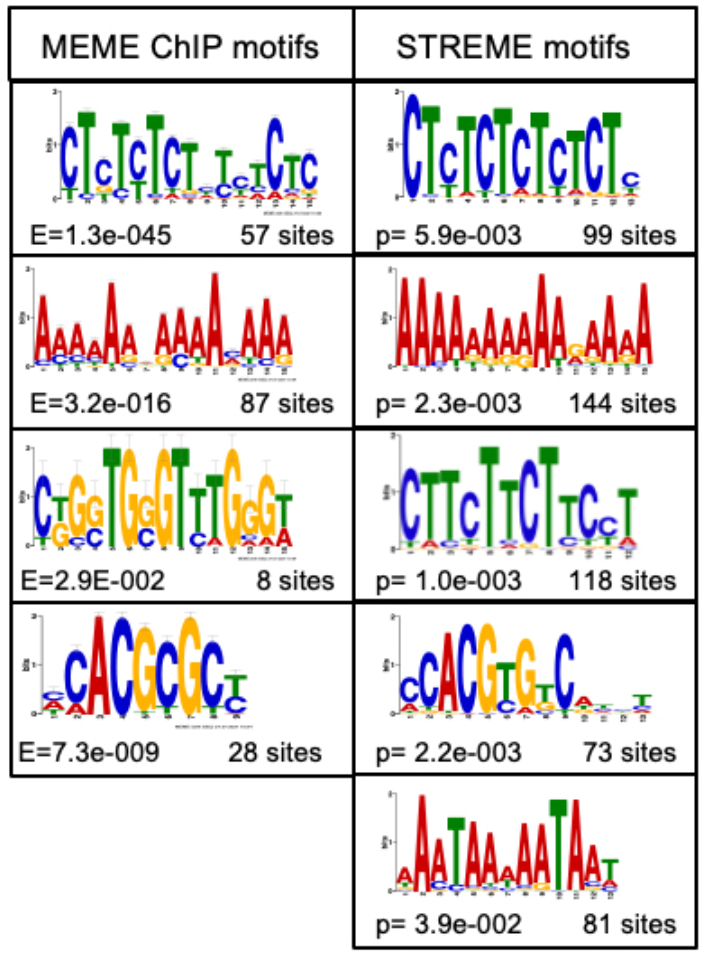
Motif discovery by MEME-ChIP and STREME. The 292 HOP2 peaks were screened by MEME-ChIP and STEME to identify conserved sequences. The E and p values, as well as the number of sites present, are shown. The CT repeat motif is similar to the binding sites of BBR/BRC transcription factors, the A-rich motif is similar to the binding sites of the REM family, and the CACGTG motif is recognized by the CAMTA family. The A-rich and GAA/CTT motif (identified by STREME) are conserved around recombination hotspots in a variety of plant species.

### HOP2 may target loci enriched for histone variants

We also conducted coimmunoprecipitation studies in conjunction with LC-MS/MS analyses to identify HOP2 binding partners, and to search for possible chromatin modifications that might play a role in HOP2 localization. It is well known that MND1 and HOP2 form a complex in vivo (Chen et al. 2004; Petukhova et al. 2005; Enomoto et al. 2006; Ploquin et al. 2007; Uanschou et al. 2013), and indeed we found a similar number of spectra for both proteins (Table 1), confirming that in our tagged line HOP2 does interact with MND1 to form a complex and carry out its functions. Several other proteins are presumed interactors based on there being few or no spectra for these in the L*er* negative control. These include the heat shock protein HSP70-3, three ribosomal subunits, CYP19-2, lipoxygenase, and ACT4, all of which are moderately abundant proteins. The significance of these enrichments is unclear.

**Table 1:** MS/MS analysis following co-immunoprecipitation. Control (Ler) and experimental (LC831) chromatin extracts from inflorescences were subjected to affinity chromatography on anti-HA paramagnetic beads. Following trypsin digestion and MS/MS analyses, the number of spectra for each identified target were recorded. The data is parsed for control samples that produced no spectra and cut off at 7 spectra for the LC831 sample, with the exception of the histone proteins listed. Post translational modifications (PTM) were also identified. K11ac represents histone H2B, acetylated at lysine 11, while K27me is histone H3 methylated at lysine 27.

| Protein | Ler spectra | LC831 spectra | PTM |
| --- | --- | --- | --- |
| HOP2 | 0 | 66 | - |
| MND1 | 0 | 59 | - |
| H2B.11 (HTB4) | 0 | 21 | K11ac |
| HSP70-3 | 0 | 18 | - |
| 40S ribosomal RPS14B | 0 | 17 | - |
| 40S ribosomal RPS5B | 0 | 14 | - |
| ACT4 | 0 | 12 | - |
| 60S ribosomal RPL8C | 0 | 8 | - |
| CYP19-2 | 0 | 8 | - |
| Lipoxygenase | 0 | 7 | - |
| H3.1 (HTR2) | 0 | 5 | K27me |
| H2A.W.7 (HTA7) | 0 | 5 | - |
| H2A.Z.9 (HTA9) | 0 | 4 | - |

It is intriguing that there also are enrichments for histone variants H2B.11 (HTB4), H3.1 (HTR2), H2A.5 (H2A.W.7), and H2A.3 (HTA.Z.9), and modifications of two of these variants. Other than the HOP2/MND1 proteins, H2B.11 (HTB4) showed the most pronounced enrichment in our tagged line and is post translationally modified by the addition of an acetyl group on LYS11. Histone H3.1 is found predominantly in transcriptionally silent regions of the genome, and the H3K27me1 modification has been associated with the maintenance of transcriptional silencing (Stroud et al. 2012; Jacob et al 2014). In a similar vein, the H2A.W variant is localized to constitutive heterochromatin and promotes chromatin condensation (Yelagandula et al. 2014; Bourguet et al., 2021), which may be coupled to the conformational changes necessary for proper chromosome segregation.

## Discussion

Our ChIP-seq experiments demonstrated that HOP2 binds along chromosome axes but not in centromeric/NOR regions, and not to the organelle genomes. With regard to the NOR regions, it has been documented that the Arabidopsis NOR regions on chromosomes 2 and 4 associate with one another and play a role in telomere clustering (Armstrong et al. 2001; Fransz et al. 2002; Pecinka et al. 2004), and that this association is independent of a number of proteins involved in pairing/recombination, including HOP2 and MND1 (Panoli et al., 2006; Stronghill et al. 2010; Da Ines et al., 2012). Thus, there may be both HOP2-dependent and HOP2-independent mechanisms of pairing. The HOP2-independent pathway may be necessary due to the sequestration of NOR sequences within the nucleolus, where they do not acquire chromatin modifications, exhibit many fewer double strand breaks, and errors are repaired by nonhomologous end joining (Sims et al. 2019).

### HOP2 exhibits preferential binding to T-DNA derived sequences

The HOP2/MND1 complex plays a role in homologous chromosome synapsis (Leu et al. 1998; Petukhova et al. 2003; Kerzendorfer et al. 2006; Vignard et al. 2007; Stronghill et al. 2010; Shi et al. 2019), and in the modulation of recombinase (DMC1/RAD51) activities (Petukhova et al. 2005; Plonquin et al. 2007; Neale and Kenney 2006). The enrichment of HOP2 binding at the *PP2A3/HOP2* locus suggests that the locus could be a bona fide pairing center, but it is well separated from both the telomere and centromere regions (*PP2A3/HOP2* at 4.5Mb; the centromere is located at ∼14Mb), where some initial associations that precede pairing occur (Hurel et al. 2018; Sepsi and Schwarzacher 2020; Aguilar and Prieto 2021 and references therein). Nevertheless, cytological analyses have shown that numerous interstitial pairing sites exist (Weiner and Kleckner 1994; Lopez et al. 2008; Higgins et al. 2012; Sepsi et al, 2017; Hurel et al. 2018), though the molecular nature of these sites is not known. As such, the *PP2A3/HOP2* locus may harbor sites involved in homology testing (see the following section). In many species, tethering of telomeres to the nuclear envelope or nucleolus, and centromere clustering occur, with subsequent movements necessary for homologous pairing (Alleva and Smolikove 2017; Link and Jantsch 2019; Zetka et al 2020; Shan et al 2021). Such movements may induce local distortions/unwinding of the DNA, particularly at A/T rich sequences. The second intron of HOP2 has a long stretch of alternating A/T that could facilitate unwinding as a prelude to the homology check.

Alternatively, the enrichment of HOP2 binding at the *PP2A3/HOP2* locus could be due to structural features of T-DNA insertions and their potential interactions. By generating wildtype and *hop2-1* haploid lines, we recently demonstrated that HOP2 is required to prevent illegitimate connections between non-homologous chromosome regions (Farahani-Tafreshi 2022), and as such, regions of the genome attempting to interact but which contain heterologous sequences would be expected to attract HOP2. There are two potential scenarios that may be relevant to our HA tagged line. First, partial homology of the two T-DNA insertions [the first being the original insertion into *HOP2* to generate the *hop2-1* mutant (Schommer et al 2003), and the second being the HA-tagged *HOP2* gene we designed to construct LC831 to enable ChIP-seq and coimmunoprecipitation studies], may be recognized as sites where perfect homology does not exist. We employed TAIL-PCR followed by sequencing to discover that the HA-tagged *HOP2* construct was mobilized into the promoter region of the starch synthase 2 gene (At3g01180). Thus, if chromosomal regions harboring the two T-DNAs attempt to pair, HOP2 may bind to exercise its role in preventing nonhomologous exchanges. Secondly, although the *hop2-1* mutation is fixed in LC831, there is the potential for a hemizygous situation, as the HA-tagged HOP2 gene construct may have inserted at multiple sites, and not all of these are present on both chromosomes. More rigorous TAIL-PCR and/or genome sequencing would be required to test this hypothesis. Importantly, a significant number of reads assigned by BowTie to the *HOP2/PP2A3* locus bear an SNP that identifies them as originating from the tagged HOP2 gene. In either of the two scenarios outlined, chromosome regions exhibiting partial homology (e.g. two T-DNAs inserted into different chromosomes) that attempt to pair would attract HOP2 and its cofactors, which would then exercise their role in preventing illegitimate exchanges.

### Does HOP2 target specific DNA sequences or chromatin motifs

HOP2 is a conserved protein that has been investigated in organisms as diverse as yeast and humans, and there exists an extensive literature on in vitro studies of HOP2 interactions with single stranded and/or double stranded DNA (Chen et al. 2004; Petukhova et al. 2005; Enomoto et al. 2006; Pezza et al. 2006; Chi et al. 2007; Pezza et al., 2007; Ploquin et al. 2007; Pezza et al. 2010; Uanschou et al. 2013; Moktan et al. 2014; Zhao et al. 2014; Kang et al. 2015; Tsubouchi et al. 2020). However, these studies have employed either viral DNA (e.g. phiX174), plasmids (pUC18), or arbitrary oligonucleotides, and to our knowledge, the potential for sequence specific binding by HOP2 has not been tested. Structural analysis of HOP2 suggests that it contains a winged helix DNA recognition motif, that is structurally similar to other sequence specific DNA binding proteins of helix-loop-helix family (Moktan et al. 2014; Zhao et al. 2014; Kang et al. 2015; Moktan and Zhou 2018). Motif analyses of our ChIP-seq data suggests that HOP2 may target one or more DNA sequences, some of which share homology with binding sites recognized by several transcription factor families. For example, the reverse complement of the (CT)^n^ sequence is similar to the canonical 5’ RGARAGRRA 3’ consensus sequence of the *BARLEY B RECOMBINANT/BASIC PENTACYSTEINE (BBR/BPC*) family that plays roles in a variety of developmental processes, and are known to recruit Polycomb repressive complexes (PRCs) that sponsor H3K27 methylation and induce chromatin compaction/silencing (Hecker et al. 2015; Mu et al. 2017; Xiao et al. 2017; Theune et al. 2019). It is possible that HOP2 binding to BBR/BPC type sequences modulates their expression to facilitate chromatin structure changes associated with meiotic chromosome compaction. In this regard, by LC-MS/MS analyses, we found that HOP2 is associated with histone H3.1 and the spectra indicate monomethylation at K27, a modification which plays roles in maintaining transcriptional silencing and genome stability in heterochromatin (Jacob et al. 2009; Dai et al., 2017; Dong et al. 2021). We also found enrichment for H2A.W and H2A.Z variants, which largely play negative roles in gene expression (Lei and Berger, 2020). The association of these three variants with HOP2 suggests that the protein could play a role in preventing recombination between repetitive DNA sequences to ensure genome integrity.

The A-rich sequence we discovered may be targeted by a subclass of the B3/REM (REproductive Meristem) transcription factors (Romanel et al. 2009; Mantegazza et al 2014; see Fig. S4). Several recent reports implicate REM genes in controlling flowering and reproductive development. REM22 is involved in specification of ovule number (Gomez et al 2018). REM16 is involved in controlling flowering time (Yu et al. 2020), and recent RNAi investigations have shown that compromising REM34/35 expression results in postmeiotic defects in both male and female gametes (Caselli et al. 2019). It has been reported that HOP2 can act as a transcription factor in mice, where it promotes the action of the osteoblast transcription factor ATF4 (Zhang et al. 2019). Whether HOP2 acts directly as a transcription factor in plants is presently unknown.

### HOP2 binding sites index potential crossover sites

One of the initial events that precedes pairing and recombination is the cleavage of DNA by SPO11 to create double strand breaks (DSBs). Choi and coworkers (Choi et al. 2018), sequenced SPO11-linked oligonucleotides and observed enrichment in gene-rich chromosome arms and depletion in heterochromatic pericentromeres, a situation that generally parallels the distribution of HOP2 binding sites. In addition, we observed that the majority of HOP2 binding sites are found in gene promoters and downstream from genes, a distribution similar to that of crossover hotspots found in Arabidopsis (Wijnker et al. 2013; Choi et al. 2013; Shilo et al. 2015; Sun et al. 2019), tomato (Demirci et al. 2017; Fuentes et al. 2020), rice (Marand et al. 2019), and maize (Pan et al. 2017). At the DNA level, motif analyses of HOP2 ChIP seq data revealed the conservation of several sequences that are identical or very similar to those found at crossover hotspots in several plant species (Choi et al. 2013, Wijnker et al. 2013; Choi et al. 2018; Pan et al. 2017). These include an A-rich repeat and a CTT repeat, a related CT repeat, and an A/T-rich sequence.

Other features of the genomic landscape associated with recombination hotspots include enrichment of histone H2A.Z, low nucleosome density, low DNA methylation and the presence of H3K4me3 (Yelina et al. 2012; Wijnker et al. 2013; Choi et al. 2013; Kianian et al. 2018; Yamada et al. 2018). Our LC-MS/MS experiments revealed an association of HOP2 with H2A.Z.9 (HTA9) but we did not examine either nucleosome density or DNA methylation. Interestingly, we also observed a significant number of spectra for H2B.11 (HTB4), some of which carry an acetyl group on K11. This modification has been reported for H2B in Arabidopsis as part of epigenome surveys (Bergmuller et al. 2007; Zhang et al. 2007; Li et al. 2022), but no literature reports exist on its potential role in meiosis. However, it has been shown that hyperacetylation of H3 is associated with altered patterns of recombination and chromosome segregation defects in Arabidopsis (Perrella et al. 2010), that a mutation in a histone deacetylase gene alters the distribution of DSBs in yeast (Mieczkowski et al. 2007), and that histone acetylation marks affect both hotspot activity and crossover resolution in mice (Getun et al. 2017). The correlation between these epigenetic marks and meiotic progression in a variety of species encourages speculation that the Arabidopsis H2B.11 variant and the acetylation of LYS11 plays a role in HOP2 biology. Significant divergence of the N-termini of Arabidopsis H2Bs (Jiang et al. 2020), and the lack of antibodies directed against the H2B.11K11ac modification currently limit further investigation. Taken together, the data are consistent with HOP2 binding to sites at or near crossover sites, possibly by recognizing a combination of genetic and epigenetic features.

## Materials and Methods

### Plant material and growth conditions

*hop2-1* seeds were kindly provided by Dr. Robert Sablowski of the John Innes Center (Schommer et al. 2003). All seeds were sown on Premier PROMIX PGX (Plant Products), and plants were propagated at 22°C in Conviron AC60 environmental chambers for 16hour light/8hour dark periods employing fluorescent lighting at ∼120μE/m^2^.

### Recombinant DNA methods

Construction of a C-terminal 3X HA tagged HOP2 gene was accomplished in two steps, engineering in restriction sites (undelined below). First, the region containing 4.7kb of upstream sequence and the HOP2 coding region was generated by PCR of Columbia DNA, using Phusion DNA polymerase (NEB) and the primers (FOR 5’C<u>TTAATTAA</u>TTAAAAGACAACAACGACCTGAATCT3’ and BACK 5’ CAT<u>GGCGCGCC</u>CTCGAGGCCTCTTTTTACCATGTTG3’), and was mobilized into the PacI and AscI sites of the binary vector pEGAD-link. A 3X HA tag-encoding region was then added to this construct by insertion of a synthetic GeneArt (Thermo-Fisher)116bp AscI-ApaLI fragment (5’ <u>GGCGCGCC</u>ATGGTTTACCCATACGATGTTCCTGACTATGCGGGCTATCCCTATGACGTCCCTGATTATG

CAGGTTCCTATCCATATGATGTTCCAGATTACGCGGTTTGA<u>GTGCAC</u>3’), which generated an in-frame fusion of the HOP2 and HA tag coding regions. The construct was mobilized into *Agrobacterium tumefaciens* GV3101 cells and used to transform *hop2-1* heterozygous plants. Second generation BASTA resistant plants were genotyped to identify *hop2-1* nulls carrying the tagged HOP2 gene.

### Genotyping

*hop2-1* mutants were genotyped by employing the wildtype primers (FOR 5’GCACTTGATAGTCTTGCTGATGCTG 3’ and BACK 5’CTCACCAATGTAATCCCTTCACG 3’) to generate a 489bp product. The forward primer was used with the T-DNA primer GABI-KATo8549 (5’ GCTTTCGCCTATAAATACGACGG 3’) to generate a PCR product of ∼340bp in samples containing an interrupted HOP2 gene.

### TAIL-PCR

TAIL-PCR was conducted essentially as described (Garrido et al. 2021), using the same arbitrary primers, but with LB1.3 (5’ ATTTTGCCGATTTCGGAAC 3’) directed towards the left T-DNA border, and EGAD BACK (5’ CCTGACTCCCTTAATTCTCCGCTCAT 3’) directed towards the right T-DNA border.

### Chromatin sample preparation and ChIP-seq

Inflorescences of LC831 and Ler were dissected into cold PBS on ice from approximately 4-5 week old plants, and all open flowers were removed. The PBS was removed and replaced with 1% formaldehyde in PBS, and vacuum infiltration and quenching was conducted as described by Yamaguchi et al. (2014). The inflorescences were washed in fresh PBS, blotted dry and either used immediately or flash frozen and stored at -70°C.

For coimmunoprecipitation, approximately 2.5 g of cross-linked inflorescences were homogenized in liquid nitrogen using a mortar and pestle. The powder was mixed with 4.5 mL Extraction Buffer [EB: 20 mM Tris-HCl pH 7.5, 150mM NaCl, 10% glycerol, 2 mM EDTA, 0.1% Igepal CA630, 1x protease inhibitor cocktail (P9599, Sigma)] and the mortar containing the frozen mixture was placed at 4°C until the mixture thawed completely. The extract was transferred into several 2 mL tubes and clarified by centrifugation in a microfuge for 10 minutes at 4°C. The clarified extracts were mixed with 50ul of EB pre-washed anti-HA coupled Protein G magnetic beads (12CA5, Thermofisher) and incubated for 2.5 hrs at 4°C on a rotator. The tubes were placed on a magnetic rack to pellet the beads and washed 4 times with EB and once more with 150 mM NaCl to remove the EB reagents. The beads, containing bound proteins, were processed for on-bead trypsin digestion and mass spectrometry analysis (LC-MS/MS) at the SPARC (SickKids Proteomics, Analytics, Robotics, & Chemical Biology Centre) Biocentre in Toronto.

For ChIP-seq and ChIP-qPCR analyses, protein-chromatin complexes were prepared from LC831 and Ler cross-linked inflorescences as previously described (Bowler et al. 2004), up to the point of chromatin shearing, which was accomplished by using micrococcal nuclease (MNase, New England Biolabs) instead of sonication.

Following the removal of EB3 (10mM Tris-HCl pH8.0, 1.7M sucrose, 0.15% Triton X-100, 2mM MgCl2, 5mM 2-mercaptoethanol, 1x protease inhibitor cocktail), the pellet was resuspended in 400ul 1x MNase buffer (50mM Tris-HCl pH 7.9, 5 mM CaCl2) and incubated at 37°C with 200 units of MNase per 100 ug of DNA. Pilot reactions were employed to determine the time of incubation that would lead to the appearance of mono/di/tri nucleosomes, as assessed by gel electrophoresis. EDTA was added to a final concentration of 10 mM, and nuclei were lysed by the addition of SDS to a final concentration of 1%. The samples were incubated on ice for 30 minutes with occasional mixing and clarified by microcentrifugation for 10 minutes at 4°C. The supernatant was diluted 10 fold with ChIP Dilution Buffer (16.7mM Tris-HCl pH8.0, 167mM NaCl, 1.1% Triton X-100, 1.2mM EDTA) and 5% was set aside as the input fraction. The remainder was incubated with equilibrated anti-HA coupled Protein G magnetic beads (12CA5, Thermofisher) for 2.5 hours at 4°C on a rotator. The tubes were placed on a magnetic rack to recover the beads, which were then washed as previously described (Yamaguchi et al. 2014). The beads were mixed with 50ul

Nuclei Lysis Buffer (NLB, 50mM Tris-HCl pH8.0, 10mM EDTA, 1% SDS) and incubated at 65°C for 30 minutes for elution of chromatin complexes. The elution was repeated once, and the pooled samples were adjusted to 0.2M NaCl in NLB and incubated at 65°C overnight to reverse the crosslinking. DNA was purified by employing a Qiaquick column as described by the manufacturer (Qiagen). In total, two pairwise LC831/L*er* control samples and two independently prepared LC831 samples were generated and outsourced to The Centre of Applied Genomics (Hospital for Sick Children, Toronto) for high-throughput sequencing where libraries were prepared and sequenced using the Illumina HiSeq2500 platform. One of the L*er* control libraries and one of the independently prepared LC831 libraries did not pass quality control tests and were excluded from the analyses. ChIP-seq sequences are available from the Sequence Read Archive (https://www.ncbi.nlm.nih.gov/sra): accession number PRJNA733468.

### Data analysis

The analysis of the ChIP-seq data was carried out using the TAIR10 reference genome. The quality of the libraries was assessed using FastQC v.11.5. Adaptor sequences were removed by employing Trim Galore v.0.0.4 and Cutadapt v.1.10. The Quality Phred score cutoff was set at 25. At least 28.8M processed reads were generated for each library. Reads were aligned to the TAIR10 genome using Bowtie2 v2.3.2 with default parameters (Langmead and Salzberg 2012). Over 91.5% of all libraries aligned to the Arabidopsis genome. Peaks were identified by MACS2 v.2.1.1 (Zhu et al. 2010) with q<0.05. Peak distribution was visualized using the Integrated Genome Browser IGB v.9.1.6 (Freese et al. 2016), and the intersection of three experimental samples was determined. Peak annotations were carried out with ChIP-peakAnno v.3.26.0 (Zhu et al. 2010), using R v.4.1.0 and the feature set of TAIR10 and TxDb.Athaliana.BioMart.plantsmart28. Peaks were classified as residing in promoter, terminator or intergenic regions, as well as in genic regions such as exons, introns, 5’ and 3’ untranslated regions (UTRs).

ChIPpeakAnno was also used to assess the distance of peaks from transcription start sites (TSS). Motif searches were conducted using STREME v.5.3.3. Source codes for bioinformatic analyses including ChIPpeakAnno are available in the project ‘git’ repository (https://bitbucket.org/nfreese/hop2-chip-seq/src/main/). Instructions for visualizing the sequencing data in IGB can also be found within the project ‘git’ repository.

For LC-MS/MS analysis, we employed Scaffold_4.11.1 with a protein threshold of 95% and a 0.2% peptide false discovery rate.

### qPCR

qPCR was performed on DNA from Ler and LC831 ChIP samples. DNA was purified from the input and bound fractions and used as temples for triplicate reactions using a QuantStudio 3 Real-time PCR system (Thermofisher Scientific) according to manufacturer’s instructions. The RTM3 gene was used as an internal control as we found this sequence was equally represented in ChIP-seq reads from both samples. Expression data was analyzed using the comparative Ct (2^−ΔΔCt^) method, where the expression values for L*er* ChIP and 831 ChIP were normalized to the RTM3 gene prior to calculating the fold enrichment of the 831 ChIP sample over the L*er* ChIP sample. We also performed qPCR on input DNA of L*er* and LC831 that was progressively digested by MNase to ascertain if rDNA sequences are present. Primer sets used for QPCR are listed in Table S1.

## Author Contributions

Conceptualization: CDR. Performed experiments: JD, MK, CDR. Software development, bioinformatics, data curation: NHF, AEL. Wrote the paper: CDR. Edited the paper: all authors. Funding acquisition: AEL, CDR. Supervision: AEL,CDR.

## Funding

This research was supported by a Natural Sciences and Engineering Research Council grant (RGPGP-2015-00071) to CDR and Dr. Clare Hasenkampf, and by grants from the National Institutes of Health (NIH NIGMS 5R01GM121927, 5R01GM103463 and 1R35GM139609 (to AEL).

## Data Availability Statement

ChIP-Seq sequences are available from the Sequence Read Archive (https://www.ncbi.nlm.nih.gov/sra) accession number PRJNA733468. Mass spectrometry data files (.raw), aligned sequencing files (.bam), and various graph and annotation files are available from CyVerse (https://data.cyverse.org/dav-anon/iplant/projects/BioViz/ChIP-Seq/PRJNA733468).

### Acknowledgments

The authors would like to thank Drs. Clare Hasenkampf, Rongmin Zhao, Sonia Gazzarrini, and Adam Mott for helpful advice and sharing of equipment. We acknowledge Nick Garrido, Selene Anton, Bruno Chue, Kimberly Bhikhari, Yisell Farahani-Tafreshi, Naden Krogen, Yuhai Cui, Ksenia Meteleva, and Lucas Corallo for lab assistance and/or technical advice.

## Conflicts of Interest

The authors declare no conflict of interest. The funders had no role in the design of the study; in the collection, analyses, or interpretation of data; in the writing of the manuscript, or in the decision to publish the results.

## Supplementary Materials

Figure S1: The HOP2pro>HOP2::3XHA construct rescues the *hop2-1* mutation.

Figure S2: Lack of HOP2 binding to NORs is not due to lack of rDNA templates.

Figure S3: Integrated genome browser export of chromosome 1.

Figure S4: Cistome identifies an enriched A-motif in REM gene promoters.

Table S1: Primers for Qpcr

**Figure S1:**
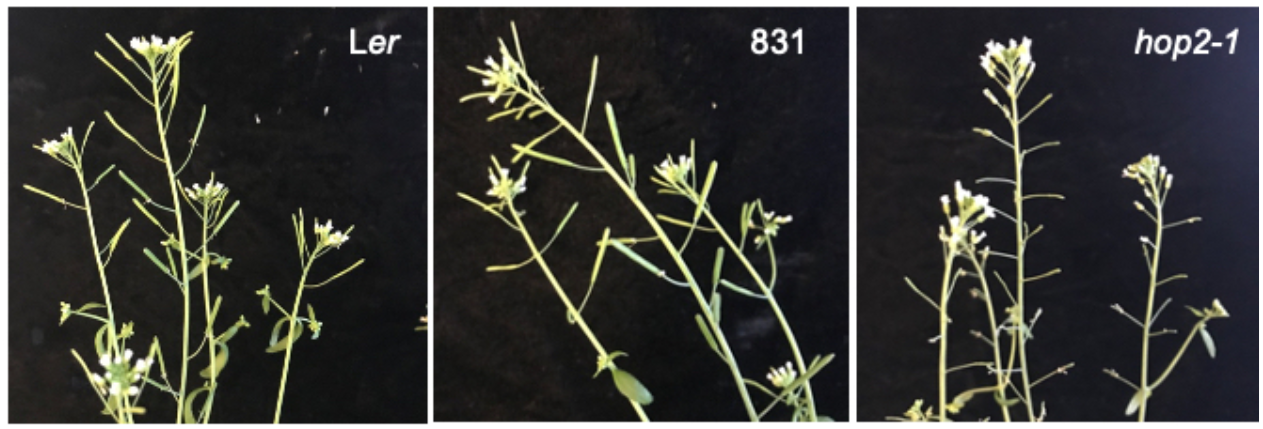
The HOP2pro>HOP2::3XHA construct rescues the *hop2-1* mutation Irftorosconcos cf the parent lino L*er* (loft}, the bop2 1 mutant (right) and the *hop2-1* mutant carrying a HOP2pro>HOP2:: 3XHA transgene (B31-centor) are shown hkxo the seventy reduced fncundty in *hop2-1* t and the mstoratKxi of A krtlity by the 831 rescue construct.

**Figure S2:**
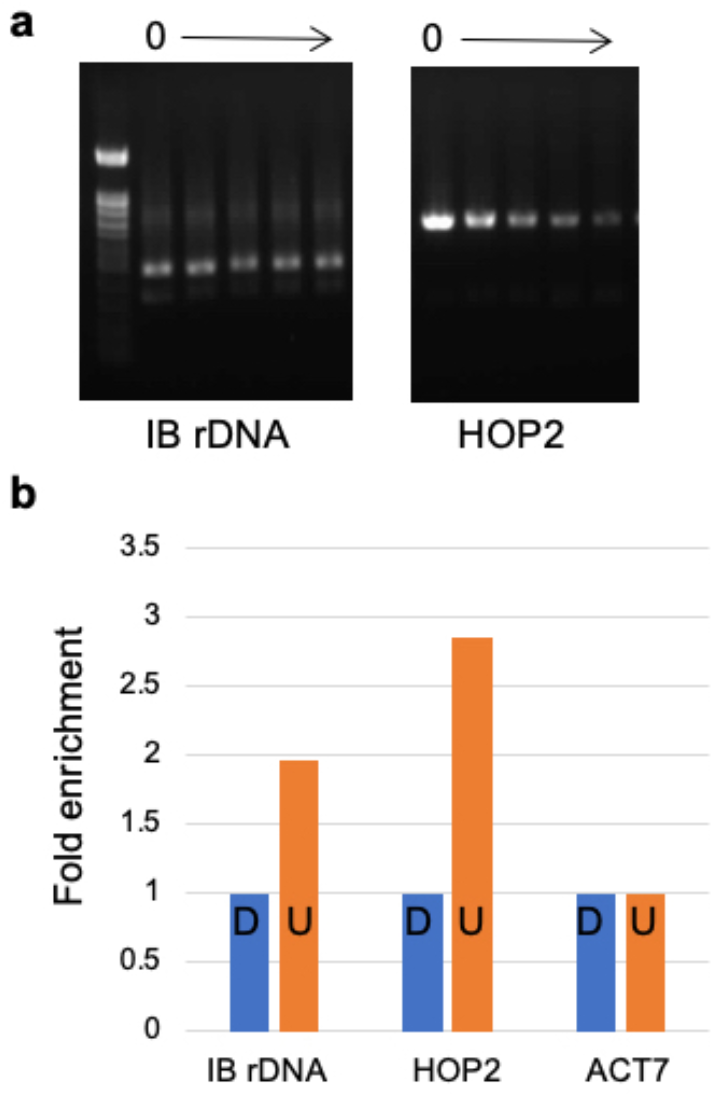
Lack of H0P2 binding to NORs Is not due to lack of rDNA templates, **a** Time course of micrococcal nuclease digestion of interval 8 rDNA and the HOP2 gene. Zero denotes no digestion. The arrow indicates progressive digestion times of 4. 8, 13. and 17 minutes. Samples were digested and the DNA purified, then templates were adjusted to the same amount of mput DNA tor subsequent PCR. **b** qPCR of undigested (U) vs. digested (D) samples (17 mnutes, producing predominantly mono and di nucleosomes) for IB rDNA and HOP2. Compared with the ACT7 control.

**Figure S3:**
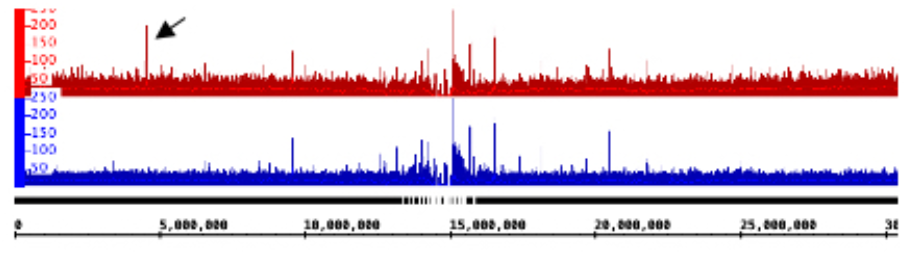
Integrated genome browser export of chromosome 1. BigWig data from an LC831 experimental sample (top, red) vs an L*er* control sample (bottom, blue). The arrow points to a significant difference in the samples at 4.5Mb, an area that harbors *the H0P2 /PP2A3* locus.

**Figure S4:**
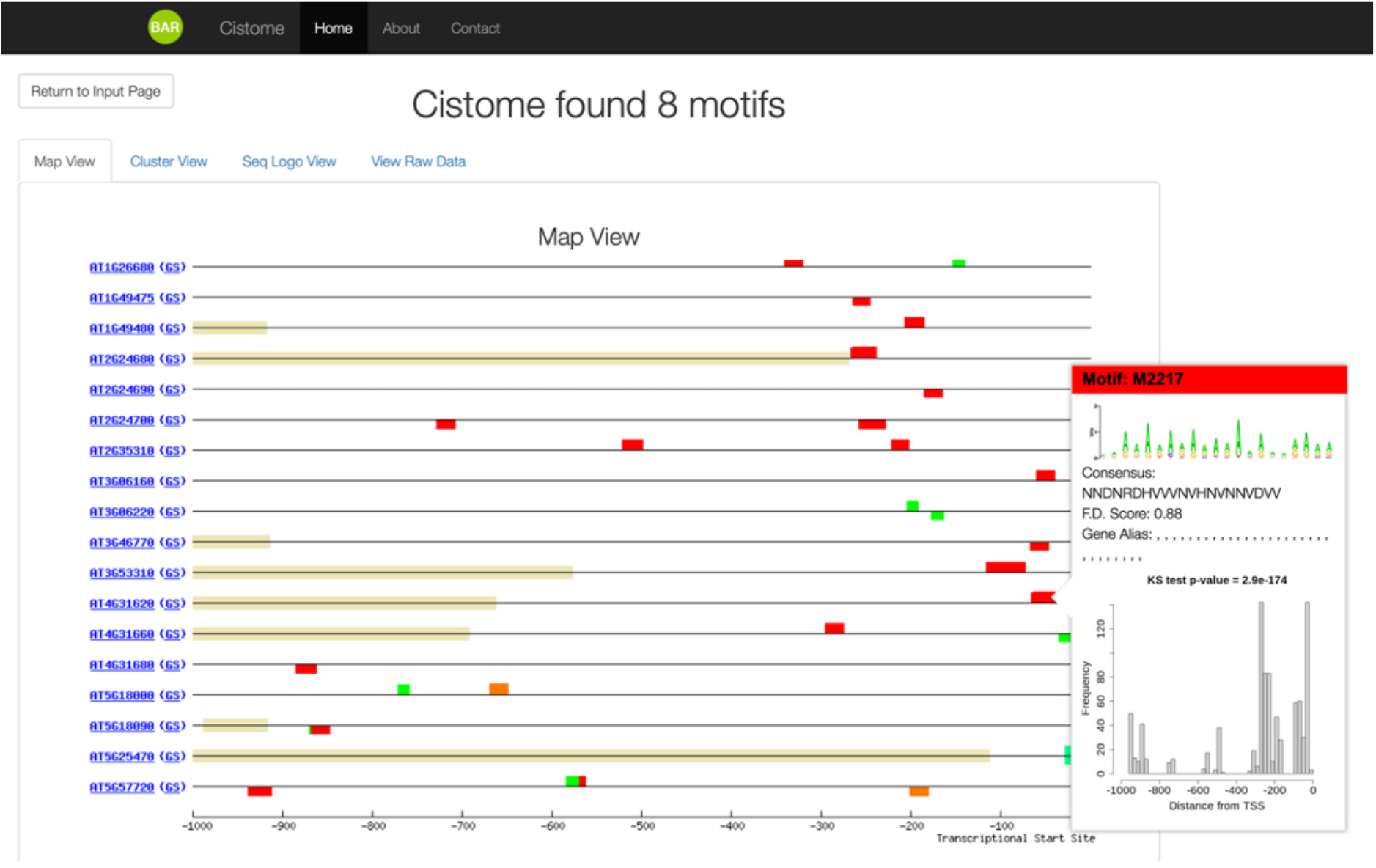
Gistome identifies an enriched A-motif. in REM gene promoters. The REM family genes (Romanel et al., 2009) were analyzed by Cistome (Austin et al. 2017), wich discovered an A-rich motif with a functional depth cutoff of 0. 88, and which was well represented in the the proximal reginons of 1000 randomly selected promoters (inset).

**Supplementary Table 1:**
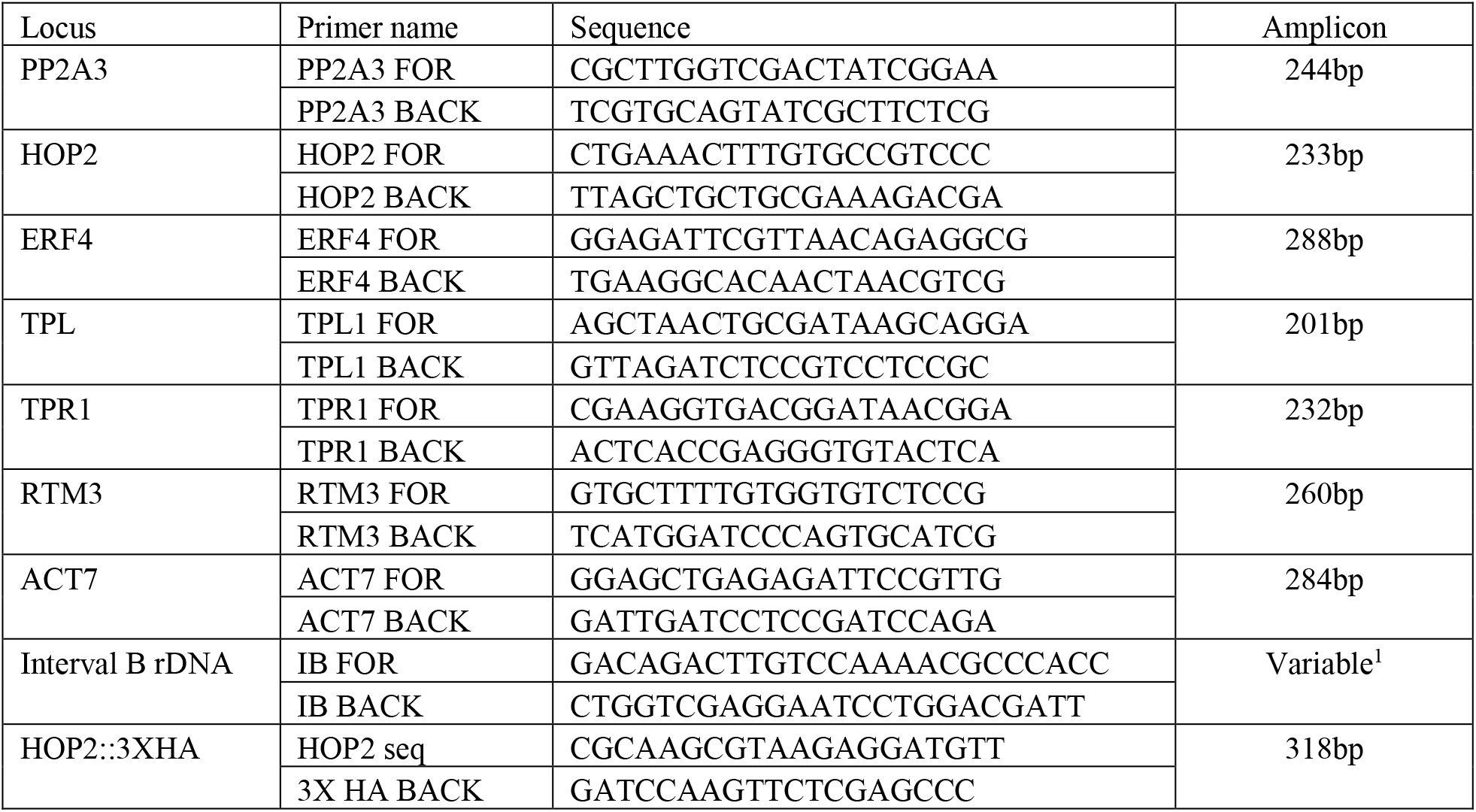
primers used for qPCR experiments

## Notes

### Competing Interest Statement

The authors have declared no competing interest.

